# Core genome MLST reveals genetic and BafA-associated phenotypic diversities in *Bartonella henselae* strains

**DOI:** 10.64898/2026.08.27.747447

**Authors:** Yoshiki Nomura, Azusa Wada, Daisuke Motooka, Masahiro Suzuki, Hidenori Kabeya, Soichi Maruyama, Shingo Sato, Kentaro Tsukamoto

## Abstract

*Bartonella henselae* is a zoonotic pathogen associated with cat-scratch disease. Although multilocus sequence typing (MLST) has been used for strain classification, its resolution for distinguishing between *B. henselae* isolates remains limited. We herein developed a *B. henselae*-specific core genome MLST (cgMLST) scheme based on whole-genome sequencing data and examined the genetic and phenotypic diversities of 80 strains derived from cats, humans, mongooses, and masked palm civets. Using the conventional MLST scheme, the 80 strains were classified into nine sequence types (STs), while cgMLST subdivided them into 72 cgSTs, demonstrating a marked improvement in discriminatory power. The cgMLST scheme comprised 1,183 core genes and showed high applicability across the 80 strains. A phylogenetic analysis revealed that ST1, which has been associated with cat-scratch disease, was further subdivided into three major clusters and two singletons, indicating high genetic heterogeneity within this ST. We also found that the *bafA* subtypes clustered in a manner that was largely consistent with the cgMLST-based phylogenetic structure, suggesting a close relationship between *bafA* variations and the genomic background of *B. henselae* strains. In a human umbilical vein endothelial cell proliferation assay, strains belonging to distinct cgSTs exhibited strain-dependent differences in proliferative capacity, which were associated with the *bafA* subtype classification. Some strains induced focal cell fragmentation and a reduced cell density at a high multiplicity of infection, indicating strain-dependent differences in endothelial cell injury. Collectively, the present results establish a high-resolution cgMLST framework for *B. henselae* and demonstrate that genetically distinct strains have diverse endothelial cell phenotypes.

## INTRODUCTION

*Bartonella henselae* causes cause cat-scratch disease (CSD), and cats are the primary reservoirs for *B. henselae* (1, 2). CSD patients generally contract the agent through scratches or bites from *B. henselae*-infected cats (3). In immunocompetent individuals, typical manifestations of CSD include papules at the site of injury, regional lymphadenopathy, fever, headache, malaise, and loss of appetite. In contrast, in immunocompromised individuals, *B. henselae* infection may lead to bacillary angiomatosis (4). *Bartonella* angiogenic factor A (BafA) was recently identified as a pathogenic factor associated with bacillary angiomatosis (5). BafA binds to vascular endothelial growth factor receptor II and activates downstream signaling pathways to induce endothelial cell proliferation and angiogenesis. This is considered to contribute to the development of vasoproliferative lesions. In our previous study (6), four sequence variants were identified in the *bafA* gene. Among these variants, the recombinant proteins BafA1 and BafA2 exhibited similar levels of angiogenic activity toward human umbilical vein endothelial cells (HUVECs). Interestingly, angiogenic activity varied among the *B. henselae* strains tested even though they carried identical *bafA* variants, suggesting the involvement of additional genomic factors. Due to the limited number of analyzed strains, the relationship between *bafA* sequence variants and the genetic backgrounds of these strains has yet to be elucidated in detail.

Numerous epidemiological studies have been conducted worldwide on the prevalence of *B. henselae* in cats (7). We previously demonstrated the prevalence of *B. henselae* in masked palm civets (*Paguma larvata*), belonging to the family Viverridae, and small Indian mongooses (*Urva auropunctata*), belonging to the family Herpestidae (8). Based on these findings, the suborder Feliformia, which comprises cat-like carnivorans, may serve as natural reservoirs for *B. henselae*. In 2001, a case of CSD caused by scratches from a masked palm civet was reported in Japan (9). Therefore, non-felid animals of the suborder Feliformia may also play a role in the transmission of *B. henselae* to humans.

Multilocus sequence typing (MLST) based on eight genes has been the most frequently used in epidemiological studies of *B. henselae*. Since the development of the MLST scheme by Iredell *et al.* (10), it has been applied in several epidemiological studies (11–19) to elucidate the genetic relationship between the sequence types (STs) of *B. henselae* strains and their hosts. To date, the MLST scheme has classified *B. henselae* strains into 41 STs, of which ST1, ST2, ST5, and ST8 were shown to correlate with CSD (11–13, 15). Arvand *et al*. suggested that ST1 strongly correlated with CSD in European countries, the United States, Israel, and Australia (11). Another study conducted in Japan reached a similar conclusion; ST1 was predominantly detected in cats and CSD patients in Japan (12). While the MLST scheme has been widely used for the epidemiological study of *B. henselae* strains, its limited genomic resolution may not fully resolve diversity within major STs, such as ST1. One major reason for its low discriminatory power is that MLST targets only a small fraction of the genome. Since whole-genome sequencing (WGS) provides access to the complete gene repertoire, a WGS-based typing scheme for *B. henselae* needs to be developed in order to obtain more detailed epidemiological and/or ecological information. Over the past two decades, next-generation sequencing techniques have been rapidly adopted worldwide for research on infectious diseases. Despite the increasing amount of bacterial WGS data, there is currently no WGS-based typing scheme for *B. henselae*.

In the present study, we aimed to develop core genome MLST (cgMLST) specific to *B. henselae* using WGS data to provide a high-resolution typing scheme. We also investigated the relationship between *bafA* sequence variants and the cgMLST-based phylogenetic classification, and specifically compared the cell proliferation-promoting abilities of *bafA1*- and *bafA2*-positive *B. henselae* strains among those classified as ST1 by conventional MLST.

## MATERIALS AND METHODS

### *B. henselae* strain collection, WGS, and WGS data retrieval

The *B. henselae* strains used in the present study are summarized in Table 1. Of these, 44 strains were cultured and sequenced to obtain the WGS data required to develop a cgMLST scheme for *B. henselae*. These strains were cultured on heart infusion agar plates containing 5% rabbit or sheep blood at 35℃ under 5% CO2 conditions for 7 days. Genomic DNA was extracted from the strains using the DNeasy Blood & Tissue Kit (Qiagen, Venlo, The Netherlands), according to the manufacturer’s instructions.

**Table 1.** *Bartonella henselae* strains used in the present study.

| Strain names | Host species | Collection<br>year | Country of<br>isolation | Isolation<br>source | Sequence data<br>source* |
| --- | --- | --- | --- | --- | --- |
| Tok.catC2-1 | *Felis catus* | 2018 | Japan | Blood | This study |
| F9-1 | *Felis catus* | 2006 | Japan | Blood | This study |
| F16-1 | *Felis catus* | 2006 | Japan | Blood | This study |
| Cat3-2022 | *Felis catus* | 2022 | Japan | Blood | This study |
| Cat29-2022 | *Felis catus* | 2022 | Japan | Blood | This study |
| Cat1-2023 | *Felis catus* | 2023 | Japan | Blood | This study |
| Cat6-2023 | *Felis catus* | 2023 | Japan | Blood | This study |
| Cat14-2023 | *Felis catus* | 2023 | Japan | Blood | This study |
| Cat15-2023 | *Felis catus* | 2023 | Japan | Blood | This study |
| Cat17-2023 | *Felis catus* | 2023 | Japan | Blood | This study |
| Cat23-2023 | *Felis catus* | 2023 | Japan | Blood | This study |
| i6-1 | *Felis catus* | 2001 | Japan | Blood | This study |
| Osa.catM35-1 | *Felis catus* | 1998 | Japan | Blood | This study |
| Osa.catM8-2 | *Felis catus* | 1998 | Japan | Blood | This study |
| Kyo.cat1 | *Felis catus* | 1998 | Japan | Blood | This study |
| Kyo.cat31 | *Felis catus* | 1998 | Japan | Blood | This study |
| Shi.cat16-2 | *Felis catus* | 1998 | Japan | Blood | This study |
| Oit.cat28 | *Felis catus* | 2008 | Japan | Blood | This study |
| Oit.cat70 | *Felis catus* | 2008 | Japan | Blood | This study |
| Oit.cat79 | *Felis catus* | 2008 | Japan | Blood | This study |
| Oit.cat98 | *Felis catus* | 2009 | Japan | Blood | This study |
| Oki.cat6 | *Felis catus* | 1998 | Japan | Blood | This study |
| Oki.cat17 | *Felis catus* | 1998 | Japan | Blood | This study |
| Oki.cat50-1 | *Felis catus* | 1998 | Japan | Blood | This study |
| IzM4 | *Homo sapiens* | 2001 | Japan | Lymph node | This study |
| 5762A | *Homo sapiens* | 2023 | Japan | Blood | This study |
| PL18 | *Paguma larvata* | 2009 | Japan | Blood | This study |
| PL186 | *Paguma larvata* | 2015 | Japan | Blood | This study |
| PL257 | *Paguma larvata* | 2025 | Japan | Blood | This study |
| HJ53 | *Urva auropunctata* | 2010 | Japan | Blood | This study |
| HJ54 | *Urva auropunctata* | 2010 | Japan | Blood | This study |
| HJ58 | *Urva auropunctata* | 2010 | Japan | Blood | This study |
| H236-1 | *Felis catus* | 2009 | Thailand | Blood | This study |
| H237-1 | *Felis catus* | 2009 | Thailand | Blood | This study |
| H238-1 | *Felis catus* | 2009 | Thailand | Blood | This study |
| H242-1 | *Felis catus* | 2009 | Thailand | Blood | This study |
| H244-1 | *Felis catus* | 2009 | Thailand | Blood | This study |
| H247-1 | *Felis catus* | 2009 | Thailand | Blood | This study |
| FR95-8724 | *Felis catus* | n/a | France | n/a | This study |
| FR95-8958 | *Felis catus* | n/a | France | Blood | This study |
| FR95-9015 | *Felis catus* | n/a | France | Blood | This study |
| U4 | *Felis catus* | 1996 | United States | Blood | This study |
| Tovarish1 | *Felis catus* | n/a | United States | n/a | This study |
| 87-66 | *Homo sapiens* | 1987 | United States | Blood | This study |
| BH1 | *Felis catus* | 2020 | Chile | Blood | GCA_030385935 |
| BH13 | *Felis catus* | 2020 | Chile | Blood | GCA_030385955 |
| BH20 | *Felis catus* | 2020 | Chile | Blood | GCA_030386185 |
| BH25 | *Felis catus* | 2020 | Chile | Blood | GCA_030386175 |
| BH27 | *Felis catus* | 2020 | Chile | Blood | GCA_030386095 |
| BH34 | *Felis catus* | 2020 | Chile | Blood | GCA_030386115 |
| BH38 | *Felis catus* | 2020 | Chile | Blood | GCA_030386215 |
| BH40 | *Felis catus* | 2020 | Chile | Blood | GCA_030386135 |
| BH45 | *Felis catus* | 2020 | Chile | Blood | GCA_030386155 |
| BH52 | *Felis catus* | 2020 | Chile | Blood | GCA_030385885 |
| BH54 | *Felis catus* | 2020 | Chile | Blood | GCA_030385975 |
| BH56 | *Felis catus* | 2020 | Chile | Blood | GCA_030386075 |
| BH58 | *Felis catus* | 2020 | Chile | Blood | GCA_030385965 |
| BH61 | *Felis catus* | 2020 | Chile | Blood | GCA_030386025 |
| BH63 | *Felis catus* | 2020 | Chile | Blood | GCA_030386055 |
| ZJBH | *Homo sapiens* | 2023 | China | Blood | GCA_029963775 |
| A233 | *Felis catus* | 1998 | Denmark | Blood | GCA_001932165 |
| A235 | *Felis catus* | 1998 | Denmark | Blood | GCA_001932225 |
| A242 | *Felis catus* | 1998 | Denmark | Blood | GCA_001932215 |
| A244 | *Felis catus* | 1998 | Denmark | Blood | GCA_001932235 |
| A20 | *Felis catus* | 1996 | France | Blood | GCA_001932075 |
| A71 | *Felis catus* | 1996 | France | Blood | GCA_001932135 |
| A74 | *Felis catus* | 1996 | France | Blood | GCA_001932085 |
| A76 | *Felis catus* | 1996 | France | Blood | GCA_001932095 |
| A112 | *Felis catus* | 1996 | France | Blood | GCA_001932145 |
| A121 | *Felis catus* | 1996 | France | Blood | GCA_001932175 |
| Marseille/URLLY-8 | *Homo sapiens* | 1996 | France | Blood | GCA_021560525 |
| FR96/BK3 | *Felis catus* | 1996 | Germany | Blood | GCA_021560385 |
| FR96/BK38 | *Felis catus* | 1996 | Germany | Blood | GCA_021560405 |
| Berlin-I | *Homo sapiens* | 1993 | Germany | Blood | GCA_021560465 |
| Zeus | *Felis catus* | 1990 | United States | Blood | GCA_000708485 |
| F1 | *Felis catus* | 1996 | United States | Blood | GCA_001932245 |
| Houston-1 | *Homo sapiens* | 1990 | United States | Blood | GCA_000046705 |
| 88-64 Oklahoma | *Homo sapiens* | 1992 | United States | Blood | GCA_021560425 |
| JK 53 | *Homo sapiens* | 1995 | United States | Blood | GCA_000708545 |
| G5436 † | *Homo sapiens* | 1995 | United States | Blood | GCA_021560445 |

n/a, not applicable
\* Numbers beginning with “GCA” refer to GenBank assembly numbers.
† This strain is synonymous with strain
Houston-1.

DNA libraries for WGS were prepared from genomic DNA with Illumina DNA Prep or Illumina DNA PCR-Free Prep (Illumina, San Diego, CA, USA), and were subsequently sequenced using the MiSeq (*n* = 25), NovaSeq 6000 (*n* = 18), and NovaSeq X Plus (*n* = 1) sequencers (Illumina, San Diego, CA, USA). The raw reads obtained were *de novo* assembled using the SKESA program (20) implemented in SeqSphere^+^ software version 10 (Ridom, Münster, Germany), and the quality of each assembly was checked with the BUSCO program (21).

The WGS data of the remaining 36 *B. henselae* strains were downloaded from “Genome” of the NCBI datasets in September 2024 (Table 1) and were used to develop and validate the *B. henselae*-specific cgMLST scheme.

### Elucidation of STs of *B. henselae* strains using the MLST scheme

The MLST scheme (10) was applied to the *B. henselae* strains examined to identify STs based on eight genes: 16S rRNA, *batR*, *ftsZ*, *gltA*, *groEL*, *nlpD*, *ribC*, and *rpoB*. If new allelic profiles were identified in the present study, new STs were assigned to the profiles and deposited in the PubMLST database (https://pubmlst.org/organisms/bartonella-henselae).

### Development of the *B. henselae*-specific cgMLST scheme

To select a set of core genes for the *B. henselae-*specific cgMLST scheme, a gene-by-gene comparison was performed on a genome-wide level using SeqSphere⁺ software version 10 (Ridom, Münster, Germany) (22). The complete genome sequence and annotation information of *B. henselae* Houston-1^T^ was selected as the “seed genome”. The other complete genome sequences of the following seven strains were used as “query genomes”: strains 88-64 Oklahoma, FR96/BK3, FR96/BK38, Berlin-I, FDAARGOS_1462, Marseille/URLLY-8, and MVT02 (Table S1).

The parameters used for cgMLST Target Definer version 1.5 in SeqSphere⁺ software (23–25) comprised the following five filters to exclude unfit genes from the seed genome: (i) a minimum length filter to discard all genes shorter than 50 bp, (ii) a start codon filter to discard all genes containing no start codon at the beginning of the gene, (iii) a stop codon filter to discard all genes containing no stop codon or more than one stop codon at the end of the gene, (iv) a homologous gene filter to discard all genes with high DNA similarity within a genome (with >90% identity and >100 bp overlap), and (v) a gene overlap filter to discard shorter genes if the two genes affected overlap >4 bp. The filtered genes were then applied to a pairwise comparison with the query genomes using BLAST version 2.2.12 (parameters used: word size 11, mismatch penalty -1, match reward 1, gap open costs 5, and gap extension costs 2). The genes of the seed genome that were shared with all query genomes with a sequence identity of ≥90% and 100% overlap were defined as a set of core genes used for the cgMLST scheme.

### Validation of the cgMLST scheme

To validate the applicability of the cgMLST scheme, the presence rates of the core genes were assessed across the 44 sequenced strains and the 30 downloaded strains, other than seed and query genomes. When at least ≥90% of the core genes were present in all strains examined (26), the typing scheme developed was deemed to be appropriate as cgMLST.

### Two phylogenetic analyses using the cgMLST scheme

Using the cgMLST scheme developed, we analyzed the WGS data of the 80 *B. henselae* strains, excluding strains FDAARGOS_1462 and MVT02 because the hosts were unknown. An allele number was automatically assigned to each core gene. Core genome STs (cgSTs) were selected based on allelic profiles, and a minimum spanning tree (MST) was constructed in SeqSphere⁺ software to visualize the genetic relationships among the strains. The phylogenetic distances among cgSTs were calculated with the pairwise ignore missing values parameter.

A core genome single nucleotide polymorphism (cgSNP) analysis was performed using the “Find Nucleotide Variants” tool in SeqSphere^+^ software. This analysis detected single nucleotide variants from the core genes in the 80 *B. henselae* strains. Subsequently, a cgSNP-based phylogenetic tree was inferred from the concatenated sequences consisting of the single nucleotide variants detected using the neighbor-joining method. To compare the discriminatory power of the cgMLST- and cgSNP-based analyses, Simpson’s diversity indices were calculated from the distribution of strains among cgSTs and cgSNP-defined clusters, respectively.

### *bafA* variants and their application to a phylogenetic analysis of *B. henselae*

To clarify the relationship between the *bafA* variants and phylogeny of the *B. henselae* strains, we extracted *bafA* nucleotide sequences from the WGS of the 80 strains by referring to a previous study (6) and identified the common *bafA* sequence region among these strains. When a new sequence variant was detected by comparing it with the four known variants (*bafA1* to *bafA4*), a new name was assigned to the sequence variant. Following this process, cgSTs constituting the MST were grouped according to the *bafA* variants.

### Cell proliferation-promoting abilities of *B. henselae* strains

We performed a proliferation assay using HUVECs to compare the cell proliferation-promoting abilities of the *B. henselae* strains. Among the culturable laboratory stock strains, one representative strain from each distinct cgST was selected, and the assay was performed on 27 of the 80 strains analyzed in the cgMLST study (Table 2). HUVECs (PromoCell, Heidelberg, Germany) were cultured in Endothelial Cell Growth Medium-2 (EGM-2; PromoCell) at 37°C under a humidified atmosphere with 5% CO2 and were used between cell passages 6 and 8 for all experiments. Regarding infection with *B. henselae* strains, HUVECs were seeded onto gelatin-coated 96-well plates at a density of 5.0 × 10^3^ cells per well in EGM-2 and were allowed to adhere to the plates for 4 h. After the adhesion period, the medium was replaced with Medium 199 (M199; Thermo Fisher Scientific, Waltham, MA) supplemented with 5% fetal bovine serum (FBS; Biowest, Nuaillé, France). *B. henselae* strains were cultured on Columbia agar plates with 5% defibrinated sheep blood at 37°C in 5% CO2 for 5–6 days. Bacterial suspensions were prepared in M199 with 5% FBS, and the bacterial cell density was adjusted based on OD600 measurements (OD600 = 1.0 was approximately equivalent to 1×10^9^ CFU/mL). HUVECs were infected at multiplicities of infection (MOIs) of 20, 100, or 500 and incubated for 72 h without a medium change at 37°C and 5% CO2. Following the incubation, infected HUVECs were stained with 10 μg/mL Rhodamine-labeled Ulex europaeus agglutinin I (UEA-I; Vector Laboratories, Newark, CA) for endothelial cell labeling and 1 μg/mL Hoechst 33342 (Nacalai Tesque, Kyoto, Japan) for nuclear staining at 37°C for 15 min. After fixation with 4% paraformaldehyde and washing with PBS, fluorescence images were acquired using a CellVoyager CQ1 Confocal Quantitative Image Cytometer (Yokogawa Electric Corporation, Tokyo, Japan). Eight replicate wells were prepared for each strain. In each well, 10 fields of view were captured using a 10× objective lens, and the total Rhodamine-UEA-I-positive cell area across the 10 fields was summed to generate one data point per well. The cell area was quantified using CellPathfinder software (Yokogawa Electric Corporation), and the proliferation rate was expressed as a fold change relative to uninfected control wells normalized to 1.0.

**Table 2.** Mapping table of *B. henselae* strains and cgST numbers identified by the cgMLST scheme.

| cgST # | ST # | Strain name | Sequence<br>variant of<br>*bafA* | HUVEC<br>assay | cgST # | ST # | Strain name | Sequence<br>variant of<br>*bafA* | HUVEC<br>assay |
| --- | --- | --- | --- | --- | --- | --- | --- | --- | --- |
| 1 | 1 | 88-64<br>Oklahoma | *bafA1* |  | 37 | 1 | F9-1 | *bafA2* |  |
| 2 | 1 | Tovarish1 | *bafA1* | ✓ | 38 | 1 | H238-1 | *bafA2* | ✓ |
| 3 | 1 | 87-66 | *bafA1* |  | 39 | 1 | H236-1 | *bafA2* | ✓ |
| 4 | 1 | Houston-1 | *bafA1* | ✓ | 40 | 1 | A20 | *bafA1* |  |
| 4 | 1 | G5436 | *bafA1* |  | 41 | 1 | HJ58 | *bafA2* |  |
| 5 | 1 | Berlin-1 | *bafA1* |  | 42 | 2 | FR95-8958 | *bafA1* |  |
| 6 | 1 | HJ54 | *bafA1* | ✓ | 43 | 4 | A235 | *bafA2* |  |
| 7 | 1 | HJ53 | *bafA1* |  | 44 | 5 | BH25 | *bafA2* |  |
| 8 | 1 | A71 | *bafA1* |  | 45 | 5 | BH1 | *bafA2* |  |
| 9 | 1 | H237-1 | *bafA1* | ✓ | 46 | 5 | BH20 | *bafA2* |  |
| 10 | 1 | Kyo.cat31 | *bafA1* | ✓ | 47 | 5 | BH63 | *bafA2* |  |
| 11 | 1 | Kyo.cat1 | *bafA1* | ✓ | 48 | 5 | BH40 | *bafA2* |  |
| 12 | 1 | Oit.cat70 | *bafA1* | ✓ | 48 | 5 | BH61 | *bafA2* |  |
| 13 | 1 | Cat23-2023 | *bafA1* | ✓ | 49 | 5 | BH27 | *bafA2* |  |
| 14 | 1 | PL186 | *bafA1* | ✓ | 50 | 5 | BH34 | *bafA2* |  |
| 15 | 1 | PL18 | *bafA1* |  | 51 | 5 | BH54 | *bafA2* |  |
| 16 | 1 | Cat29-2022 | *bafA1* | ✓ | 52 | 5 | BH52 | *bafA2* |  |
| 17 | 1 | Oit.cat98 | *bafA1* | ✓ | 53 | 5 | BH38 | *bafA2* |  |
| 18 | 1 | PL257 | *bafA1* | ✓ | 54 | 5 | FR96/BK38 | *bafA2* |  |
| 19 | 1 | BH13 | *bafA1* |  | 55 | 5 | BH58 | *bafA2* |  |
| 20 | 1 | F1 | *bafA1* |  | 56 | 5 | BH56 | *bafA2* |  |
| 21 | 1 | ZJBH | *bafA2* |  | 57 | 5 | JK 53 | *bafA2* |  |
| 22 | 1 | Shi.cat16-2 | *bafA2* |  | 58 | 5 | Zeus | *bafA2* |  |
| 23 | 1 | IZM4 | *bafA2* |  | 59 | 6 | BH45 | *bafA3* |  |
| 24 | 1 | i6-1 | *bafA2* |  | 60 | 6 | U4 | *bafA3* | ✓ |
| 25 | 1 | Oki.cat50-1 | *bafA2* |  | 61 | 6 | A233 | *bafA3* |  |
| 26 | 1 | Oit.cat28 | *bafA2* |  | 62 | 6 | Marseille/U<br>RLLY-8 | *bafA3* |  |
| 27 | 1 | Oit.cat79 | *bafA2* | ✓ | 63 | 7 | FR95-8724 | *bafA4* | ✓ |
| 28 | 1 | Cat6-2023 | *bafA2* | ✓ | 64 | 7 | A242 | *bafA4* |  |
| 29 | 1 | Osa.catM35-1 | *bafA2* | ✓ | 65 | 7 | A244 | *bafA4* |  |
| 30 | 1 | Osa.catM8-2 | *bafA2* | ✓ | 66 | 7 | FR96/BK3 | *bafA4* |  |
| 31 | 1 | Cat3-2022 | *bafA2* | ✓ | 67 | 8 | FR95-9015 | *bafA2m* | ✓ |
| 32 | 1 | Oki.cat17 | *bafA2* | ✓ | 68 | 8 | A74 | *bafA2m* |  |
| 33 | 1 | Tok.catC2-1 | *bafA2* | ✓ | 68 | 8 | A76 | *bafA2m* |  |
| 34 | 1 | Cat1-2023 | *bafA2* | ✓ | 69 | 11 | A112 | *bafA3* |  |
| 34 | 1 | Cat14-2023 | *bafA2* |  | 70 | 11 | A121 | *bafA3* |  |
| 34 | 1 | Cat15-2023 | *bafA2* |  | 71 | 42* | Oki.cat6 | *bafA3* | ✓ |
| 34 | 1 | Cat17-2023 | *bafA2* |  | 72 | 42* | H242-1 | *bafA3* | ✓ |
| 35 | 1 | 5762A | *bafA2* |  | 72 | 42* | H244-1 | *bafA3* |  |
| 36 | 1 | F16-1 | *bafA2* |  | 72 | 42* | H247-1 | *bafA3* |  |

\* ST42 was newly identified in this study.

### Statistical analysis

Statistical analyses were performed using GraphPad Prism 10 software (GraphPad Software, La Jolla, CA). A two-way ANOVA followed by Dunnett’s multiple-comparisons test was used to compare experimental groups with uninfected controls, and results are presented as the mean ± standard deviation. Regarding heatmap visualization, log2-transformed cell proliferation values were plotted using the same software. To evaluate the effects of the *bafA* subtype and MOI on HUVEC proliferation, analyses were restricted to *bafA1*- and *bafA2*-positive strains because sufficient sample sizes were available only for these two subtypes. Relative cell numbers were normalized to the uninfected control, set to 1.0, log2-transformed, and analyzed by a two-way ANOVA followed by Sidak’s multiple-comparisons test.

### Data availability

The raw reads of all sequenced *B. henselae* strains (*n* = 44) were deposited in the Sequence Read Archive of the DNA Data Bank of Japan and are available under BioProject accession numbers PRJDB17373 and PRJDB40423.

## RESULTS

### Genome sequencing and MLST classification

The *de novo* assembly process resulted in coverage ranging from 39- to 235-fold per strain. The mean number of assemblies was 150 (*n* = 25) on MiSeq and 179 (*n* = 19) on NovaSeq sequencers. Assembly completeness exceeded 95% for all strains. The 80 *B. henselae* strains were classified into nine STs: ST1, ST2, ST4, ST5, ST6, ST7, ST8, ST11, and ST42 (Table 2). Cat-derived strains (*n* = 64) were classified into nine STs as ST1 (*n* = 31), ST2 (*n* = 1), ST4 (*n* = 1), ST5 (*n* = 15), ST6 (*n* = 3), ST7 (*n* = 4), ST8 (*n* = 3), ST11 (*n* = 2), and a novel ST42 (*n* = 4). CSD patient-derived strains (*n* = 10) were divided into ST1 (*n* = 8), ST5 (*n* = 1), and ST6 (*n* = 1). Six strains from three mongooses and three masked palm civets were classified as ST1.

### Development, validation, and phylogenetic analysis of the cgMLST scheme

A total of 1,599 genes were extracted from the seed genome as core gene candidates. Following gene filtration, 416 genes were excluded and the remaining 1,183 genes were defined as core genes. The applicability of the cgMLST scheme was then validated based on the presence rates of the defined core genes. Presence rates ranged from 93.2% for strain Oki.cat6 to 100% for strain G5436, with a mean of 98.0 ± 1.6% SD (Table S2).

In the MST analysis, the 80 strains were classified into 72 cgSTs (Figure 1A and Table 2). Simpson’s diversity index for cgMLST indicated a discriminatory power of 0.999 (confidence interval: 0.998−1.000). The number of cgSTs identified within each ST was as follows: 41 for ST1 (*n* = 45), 15 for ST5 (*n* = 16), four for ST6 (*n* = 4), four for ST7 (*n* = 4), two for ST8 (*n* = 3), two for ST11 (*n* = 2), and two for ST42 (*n* = 4). To define a cut-off for clustering cgSTs, we evaluated the number of loci that differed between cgSTs and used a cut-off of 90 loci, which grouped cgSTs in a manner that broadly corresponded to the MLST-defined STs for ST5, ST7, and ST8. Accordingly, cgSTs differing by ≤90 core genome loci were grouped into the same cluster. The cgSTs of ST5, ST7, and ST8 each formed a distinct cluster, whereas those of ST6, ST11, and ST42 were grouped into a single cluster. The cgSTs of ST2 and ST4 were maintained as singletons because of the presence of one strain per one ST. The cgSTs comprising ST1 were classified into three distinct clusters (A–C) and two singletons (α and β). ST1-cluster A included *B. henselae* strains from cats and CSD patients in Japan, Thailand, Chile, France, Germany, and the United States, as well as from non-felid Feliformia, such as mongooses and masked palm civets in Japan. ST1-singleton α included a *B. henselae* strain isolated from a cat in France. ST1-clusters B and C and singleton β comprised *B. henselae* strains from cats and CSD patients in Asian countries: Japan and China for cluster B, Thailand for cluster C, and Japan for singleton β.

**Figure 1.**
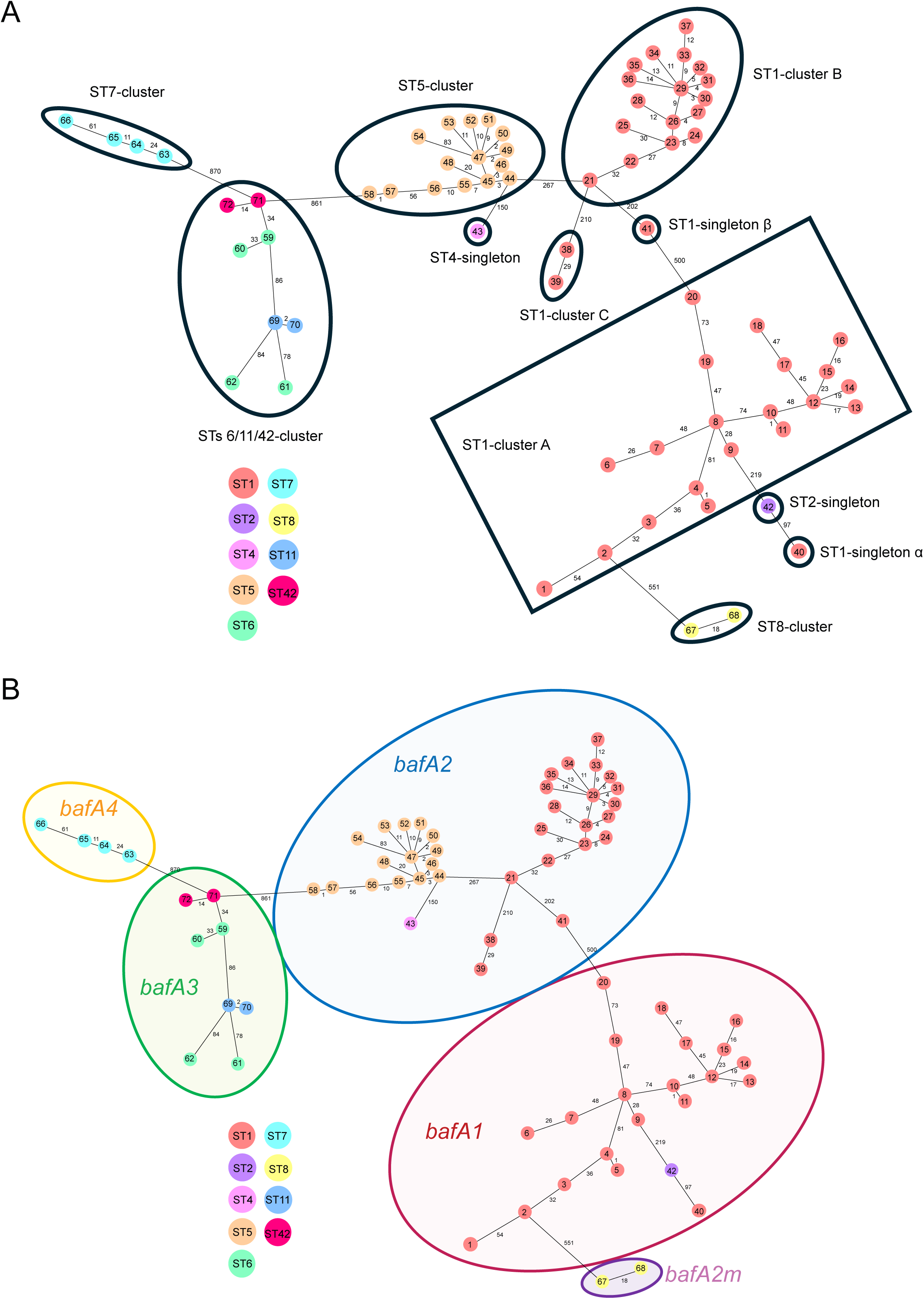
(A) A minimum spanning tree (MST) based on the allelic profiles of 1,183 core genes from 80 *B. henselae* strains. The MST was visualized using Ridom SeqSphere+ software with pairwise comparisons ignoring missing values. The numbers inside circles indicate cgSTs and circle colors indicate STs. The numbers on the connecting lines correspond to the number of core genes that differed between cgSTs. cgSTs differing by 90 or fewer core genes were grouped into the same cluster. Clusters containing multiple cgSTs and singletons containing a single cgST are enclosed by thick black lines. (B) Genetic relationship between *bafA* sequence variants and cgSTs. cgSTs were grouped according to *bafA* sequence variants: *bafA1*, *bafA2*, *bafA2m*, *bafA3*, and *bafA4*.

A total of 10,721 SNPs were detected from the multiple alignments of the core genes and were used to calculate a phylogenetic distance matrix. A phylogenetic tree based on the cgSNP schemes classified the 80 *B. henselae* strains into seven clusters and four singletons, consistent with those identified by the MST (Figure S1). Simpson’s diversity index for cgSNPs indicated a discriminatory power of 0.999 (confidence interval: 0.998−1.000).

### Genetic relationship between *bafA* sequence variants and cgSTs of *B. henselae* strains

Comparisons of *bafA* sequences among the 80 *B. henselae* strains identified a 1,218-bp region (base positions 1 to 1,218 in the gene) that was shared across all strains and may be used for *bafA* subtype comparisons. Subtyping based on this region identified a new *bafA* sequence variant in strains A74, A76, and FR95-9015. This variant shared 73.0% identity (880/1,218bp) with *bafA1*, 96.1% identity (1,170/1,218bp) with *bafA2*, 74.9% identity (912/1,218bp) with *bafA3*, and 82.5% identity (1,005/1,218bp) with *bafA4*. Therefore, the new sequence variant was designated as a *bafA2* mutation (*bafA2m*) because its homology with *bafA2* was the highest. The *bafA* sequences of the 80 strains were classified into the following five groups: 23 strains with *bafA1*, 40 with *bafA2*, three with *bafA2m*, 10 with *bafA3*, and four with *bafA4* (Table 2). The 72 cgSTs in the MST were then regrouped according to the *bafA* sequence variant. This classification formed five groups in the MST, each corresponding to one *bafA* variant category (Figure 1B).

### Differences in HUVEC proliferation abilities by *bafA* variants

HUVEC proliferation was assessed after infections with representative *B. henselae* strains selected from distinct cgSTs at MOIs of 20, 100, and 500 (Figure 2A). At MOIs of 20 and 100, all strains, except for FR95-9015 (cgST67) and Oki.cat6 (cgST71), showed significantly higher proliferation than the uninfected controls. Among these, Houston-1 (cgST4), HJ54 (cgST6), H237-1 (cgST9), Oit.cat98 (cgST17), and H238-1 (cgST38) increased the proliferation ratio by more than 1.5-fold, whereas the remaining strains showed more modest increases of approximately 1.2-fold. At an MOI of 500, the proliferative effects of most strains decreased. Osa.catM8-2 (cgST30), Cat1-2023 (cgST34), H236-1 (cgST39), and Oki.cat6 (cgST71) exhibited significantly lower cell numbers than the uninfected controls.

**Figure 2.**
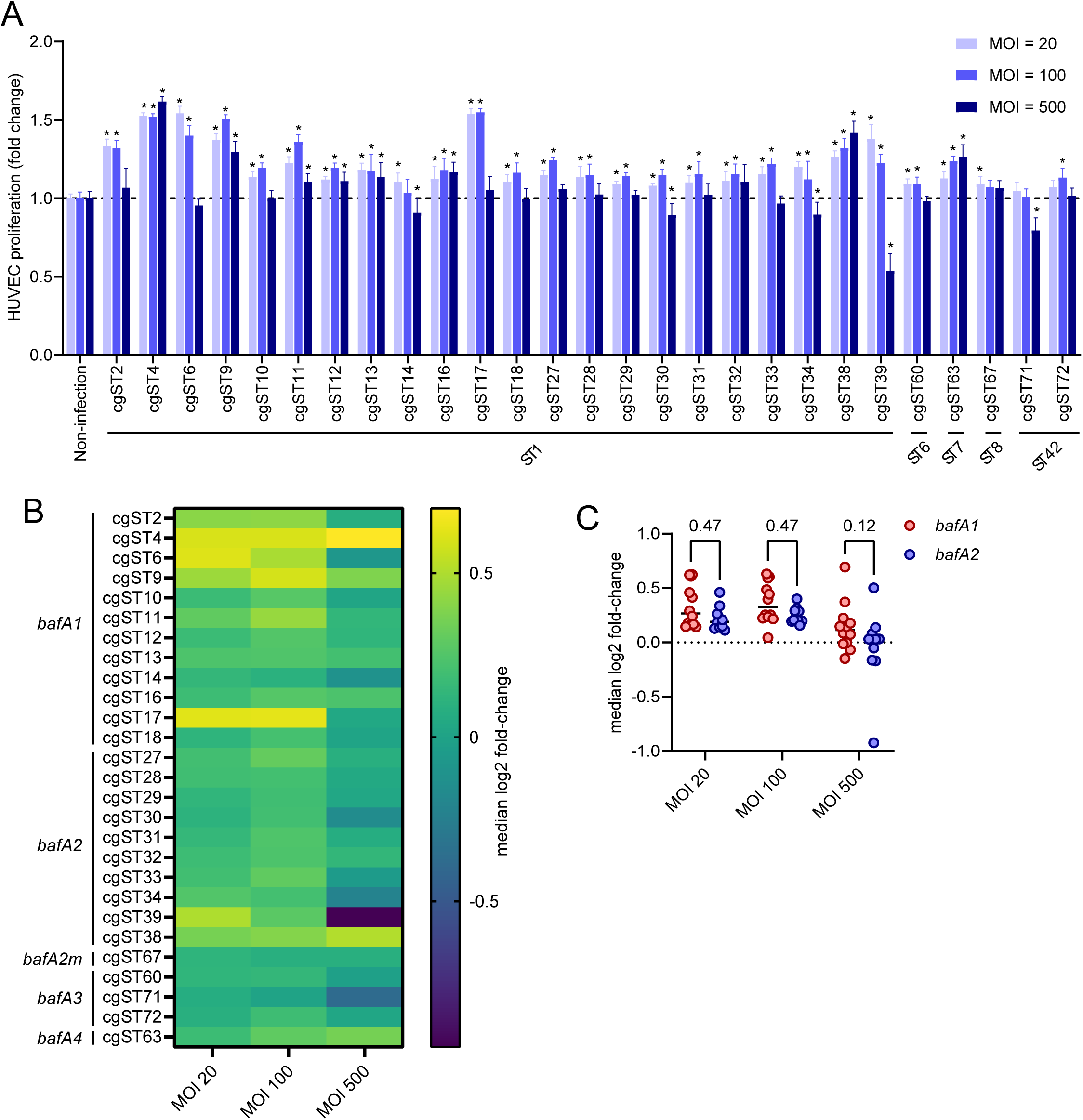
Quantitative comparison of HUVEC proliferation induced by *B. henselae* strains. (A) Fold change in HUVEC proliferation from uninfected controls following infection with representative cgST-defined strains at MOIs of 20, 100, and 500. Proliferation was quantified by summing the total Rhodamine-UEA-I-stained endothelial cell area across 10 fields per well. Each bar represents the mean ± SD of 8 wells (technical replicates). Asterisks indicate significant differences from the uninfected control (*P* <0.05). (B) Heatmap of median log2-transformed fold changes in HUVEC proliferation generated using GraphPad Prism 10. (C) Comparison of HUVEC proliferation induced by *bafA1*- and *bafA2*-positive strains at different MOIs. HUVEC proliferation values were normalized to the uninfected control and expressed as a log2 fold change. Each dot represents an independent strain. The horizontal dotted line indicates the uninfected control level (log2 fold change = 0). The significance of differences was assessed by a two-way ANOVA followed by Sidak’s multiple comparisons test; adjusted *P* values are shown above each pairwise comparison.

A heatmap analysis revealed distinct proliferative patterns associated with *bafA* sequence variants (Figure 2B). The *bafA2m*-positive strain FR95-9015 (cgST67) exhibited no detectable proliferative activity. A sequence analysis identified a frameshift mutation in the *bafA*-coding region of this strain, introducing a premature stop codon.

Since only three strains carried *bafA3* and only one strain carried *bafA4*, statistical comparisons were restricted to the *bafA1*- and *bafA2*-positive groups (Figure 2C). A two-way ANOVA of log2-transformed relative cell numbers showed the significant main effects of MOI (*P* value < 0.001) and *bafA* subtype (*P* value = 0.008), but no significant interaction between the two factors (*P* value = 0.82). Sidak’s post hoc multiple-comparisons test detected no significant difference between the *bafA1*- and *bafA2*-positive strains at any MOI tested.

Representative fluorescence images of HUVECs infected with the three strains, Houston-1 (cgST4), PL186 (cgST14), and H236-1 (cgST39), at each MOI are shown in Figure 3. In the merged images, rhodamine-UEA-I-stained endothelial cell regions (red) and Hoechst 33342-stained nuclei (cyan) revealed discrete bacterial aggregates, and their abundance increased with the MOI. Houston-1 (cgST4) infection promoted HUVEC proliferation, even at an MOI of 500. In contrast, PL186 (cgST14) infection did not consistently promote cell proliferation at any of the tested MOIs. H236-1 (cgST39) increased HUVEC proliferation at an MOI of 20, but showed a reduced cell density and focal cell fragmentation around bacterial aggregates at MOIs of 100 and 500. In H236-1-infected cultures, regions with sparse bacterial aggregates retained a higher cell density, whereas areas adjacent to bacterial aggregates showed a reduced cell density and fragmentation. Cultures with Houston-1 (cgST4) and PL186 (cgST14) showed only limited cell loss and fragmentation in the vicinity of bacterial aggregates.

**Figure 3.**
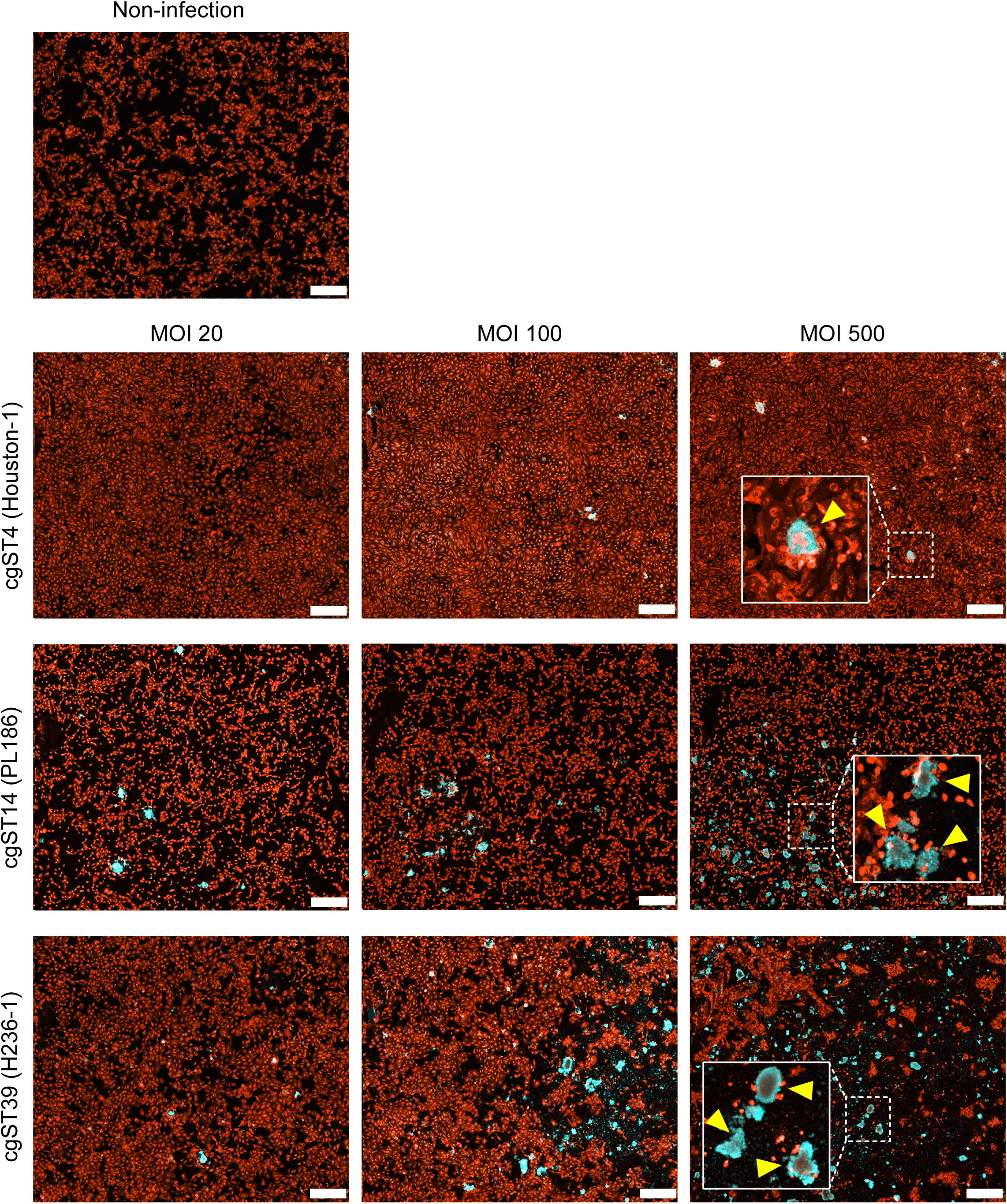
Representative fluorescence images showing strain- and MOI-dependent differences in HUVEC responses to *B. henselae* infection. HUVECs were either uninfected or infected with Houston-1 (*bafA1*, cgST4, ST1-cluster A), PL186 (*bafA1*, cgST14, ST1-cluster A), or H236-1 (*bafA2*, cgST39, ST1-cluster C) at MOIs of 20, 100, and 500. Endothelial cell regions were stained with rhodamine-UEA-I (red), while nuclei were stained with Hoechst 33342 (cyan). White boxes indicate the regions shown at a higher magnification. Arrowheads indicate bacterial aggregates. Scale bars, 500 μm.

## DISCUSSION

We herein developed a *B. henselae*-specific cgMLST scheme and applied it to 80 strains from eight countries. The conventional MLST scheme classified the 80 strains into nine STs, while the cgMLST scheme subdivided them into 72 cgSTs, demonstrating a markedly higher resolution for strain discrimination. According to a previous study (26), putative core genes found in at least 90% of the genomes examined have been defined as core genes by CoreCruncher, a Python-based program. In the present study, core gene presence rates ranged from 93.2 to 100%, indicating that the cgMLST scheme developed captured most of the target loci across the analyzed strains. These results support the broad applicability of this scheme to *B. henselae*.

Conventional MLST has been utilized for more than two decades to investigate the genetic diversity of *B. henselae* strains, with 42 STs being deposited in the MLST database by April 2026 (27). Among them, ST1 has been repeatedly linked to CSD in multiple regions, including Australia, Europe, the United States, Israel, and Japan (11, 12, 15). In the present study, ST1 accounted for more than half of the analyzed strains (45 out of the 80 strains) and was further subdivided into 41 cgSTs, indicating the high genetic diversity of *B. henselae* strains belonging to ST1.

Using a cut-off of 90 loci, cgSTs were grouped into seven clusters and four singletons, and this structure broadly agreed with the classification obtained by the cgSNP analysis. Identical Simpson’s diversity indices for the cgMLST and cgSNP schemes indicate that the two approaches have provided equivalent discriminatory power for this dataset. However, cgMLST has the practical advantage of assigning strains with identical allelic profiles to the same cgST, which may facilitate strain matching across different studies and comparisons of strains from CSD patients and host animals.

Within ST1, the cgMLST scheme resolved three clusters (A–C) and two singletons. ST1-cluster A included strains from CSD patients in the United States and Germany and cats in Japan, Thailand, Chile, the United States, and France, and strains from mongooses and masked palm civets in Japan. Notably, this cluster included four human strains from the United States, which were assigned to three cgSTs (cgSTs 1, 3, and 4), and one human strain from Germany, which was assigned to cgST5. Among these strains, Houston-1ᵀ and G5436 exhibited identical allelic profiles (cgST4). These two strains are considered to be synonymous strains, supporting the high discriminating power of the cgMLST scheme. In contrast, ST1-cluster B predominantly contained strains from cats in Japan, together with strains from Japanese and Chinese CSD patients, whereas no human isolates from Europe or the Americas were included in this cluster. These geographic patterns suggest that the population structure within ST1 differs among regions and also that ST1-cluster B may contribute differently to human infections in Asia and other regions. Additional sampling of human isolates, particularly from Asian countries, is required to further investigate this hypothesis.

The clustering patterns of ST6, ST11, and ST42 are also noteworthy. While cgMLST grouped their cgSTs (cgSTs 59–62 and 69–72) into a single cluster, cgSTs 59–62 (ST6) were separated into two populations within this cluster, with cgSTs 69 and 70 (ST11) being positioned between them. This arrangement suggests that the phylogenetic relationships among these STs are more complex than those implied by their MLST designations. According to previous studies (11, 13, 15, 16), ST6 was found to be distributed worldwide, and further isolates will be necessary to clarify the diversity of the ST6-related group.

The HUVEC proliferation assay also showed that endothelial responses markedly differed among the strains. These differences were the most apparent at lower MOIs, whereas at higher bacterial burdens, the proliferative effect was attenuated in many strains and cell numbers decreased below control levels in some cases. Therefore, the endothelial phenotype associated with *B. henselae* infection appears to depend not only on the strain background, but also on the bacterial load. Imaging results support this interpretation because focal cell fragmentation and a reduced cell density were observed near bacterial aggregates in H236-1 (cgST39)-infected cultures, whereas only limited cell fragmentation and cell losses were noted in cultures infected with Houston-1 (cgST4) and PL186 (cgST14).

The relationship between the proliferation phenotype and *bafA* subtype supports a genetic contribution to this functional diversity. The two-way ANOVA revealed the significant main effects of both MOI and *bafA* subtype, whereas no significant interaction was detected, suggesting that the effect of *bafA* subtype on HUVEC proliferation was broadly consistent across the MOI range tested. Many *bafA1*-positive strains induced HUVEC proliferation more strongly than *bafA2*-positive strains; however, post hoc comparisons within each MOI did not detect a significant difference between the two subtypes, indicating that the subtype-associated effect was modest at the individual MOI level. The *bafA2m*-positive strain FR95-9015 (cgST67) exerted no proliferative effect, and its frameshift mutation may have rendered it a loss-of-function allele. Moreover, the result showing that strains carrying the same *bafA* subtype still differed in their effects on HUVECs indicates that *bafA* sequence variations alone do not fully explain the phenotypic diversity observed. Collectively, these results suggest that additional bacterial or host factors modulate the proliferative response, and also that further experiments with larger sample sizes, additional strains, or mechanistic assays will be needed to clarify the contribution of *bafA* variations to HUVEC proliferation. Since statistical comparisons were restricted to *bafA1* and *bafA2* due to the limited sample sizes for the other subtypes, the present results need to be interpreted as subtype-specific rather than generalizable to all *bafA* variants. This point is particularly important for interpreting high-MOI phenotypes. In several strains, a reduced cell density and focal fragmentation were observed along with bacterial aggregation, suggesting that infection outcomes reflect a balance between proliferation-promoting activity and cell-damaging effects. Previous studies demonstrated that the VirB/VirD4 type IV secretion system (T4SS) and translocated *Bartonella* effector proteins modulated endothelial cell behavior and may contribute to both survival-related and cytotoxic phenotypes (28, 29). The present results are consistent with a model in which BafA-driven proliferative signaling is superimposed on additional strain-dependent activities, including contact-associated or T4SS-related effects, which become more evident at high bacterial burdens (30, 31). Accordingly, the strain-dependent endothelial phenotypes described herein may be attributed to multiple bacterial factors rather than to the *bafA* subtype alone. Differences in the expression or activity of T4SS effectors, as well as uncharacterized secreted or surface-associated molecules, may contribute to the strain-dependent differences observed in endothelial cell proliferation and injury among strains. Therefore, defining these factors is important for understanding how genomic diversity translates into differences in pathogenic potential.

Overall, the newly developed cgMLST scheme provides a high-resolution framework for characterizing the high genetic and phenotypic diversities of *B. henselae* beyond those revealed by conventional MLST. This scheme enables the discrimination of closely related strains and links phylogenetic relationships with endothelial phenotypes. This framework may serve as a valuable tool for comparative genomic studies, source attribution between human and animal isolates, and future investigations aimed at identifying the bacterial factors contributing to strain-dependent pathogenic diversity.

## ACKNOWLEDGMENTS

The authors would like to express their sincere gratitude to Professor Emeritus Bruno B. Chomel, University of California, Davis, for generously providing the *Bartonella henselae* strains used in this study. We acknowledge the NGS core facility at the Research Institute for Microbial Diseases of The University of Osaka for the sequencing and data analyses. We also thank Xingyan Ma and Hongyuan Jia for their technical assistance with the cell-based analysis. This work was supported by the Japan Agency for Medical Research and Development (AMED) Japan Program for Infectious Diseases Research and Infrastructure under grant number JP23wm0325058 (to K.T.); the Japan Society for the Promotion of Science (JSPS) KAKENHI, JP22K070602 (to K.T.); the Grant for Joint Research Project of the Research Institute for Microbial Diseases, The University of Osaka, JRPRIMD25C2 (to S.S)

## AUTHOR CONTRIBUTIONS

Y.N. performed the laboratory work, whole-genome sequencing, and data analyses and wrote the draft. A.W. managed the *B. henselae* strains and supported laboratory work by Y.N. D.M. and M.S. performed the whole-genome sequence analysis. H.K. and S.M. supervised the research and revised the manuscript. S.S. and K.T. designed the study plan, analyzed and curated data, and revised the manuscript.

