## Supplementary materials for "Core genome MLST reveals genetic and BafA-associated phenotypic diversities in *Bartonella henselae* strains"

Table S1. *Bartonella henselae* strains used to define the core gene set for the cgMLST scheme

| Purpose of use | Strain name | Country of isolation | Host species | Genome size | ST # | Refseq # |
|---|---|---|---|---|---|---|
| Seed | Houston-1 | United States | *Homo sapiens* | 1,931,047 bp | 1 | NC_005956 |
| Query | 88-64 Oklahoma | United States | *Homo sapiens* | 1,969,298 bp | 1 | NZ_CP072899 |
| Query | FR96/BK3 | Germany | *Felis catus* | 1,935,288 bp | 7 | NZ_CP072897 |
| Query | FR96/BK38 | Germany | *Felis catus* | 1,944,393 bp | 5 | NZ_CP072898 |
| Query | Berlin-I | Germany | *Homo sapiens* | 1,931,655 bp | 1 | NZ_CP072901 |
| Query | FDAARGOS_1462 | Germany | Unknown | 1,946,727 bp | 1 | NZ_CP082885 |
| Query | Marseille/URLLY-8 | France | *Homo sapiens* | 1,906,759 bp | 6 | NZ_CP072904 |
| Query | MVT02 | Unknown | Unknown | 1,905,383 bp | 9 | NZ_LN879429 |

Table S2. Presence rates of core genes among *Bartonella henselae* strains

| Strain name | Presence rate of core genes | Strain name | Presence rate of core genes |
|---|---|---|---|
| Oki.cat6 | 93.2 | JK 53 | 98.3 |
| U4 | 94.0 | Oit.cat28 | 98.3 |
| H244-1 | 94.1 | Oki.cat17 | 98.3 |
| H247-1 | 94.3 | Tok.catC2-1 | 98.3 |
| BH45 | 94.4 | Oit.cat79 | 98.4 |
| H242-1 | 94.4 | HJ58 | 98.5 |
| A242 | 94.9 | IZM4 | 98.5 |
| A244 | 94.9 | Oki.cat50-1 | 98.5 |
| FR95-8724 | 94.9 | A74 | 98.6 |
| BH25 | 95.5 | F16-1 | 98.6 |
| BH34 | 96.2 | H236-1 | 98.6 |
| A112 | 96.6 | H238-1 | 98.6 |
| A121 | 96.6 | i6-1 | 98.6 |
| A233 | 96.6 | Osa.catM8-2 | 98.6 |
| BH61 | 97.0 | 5762A | 98.7 |
| BH63 | 97.5 | F9-1 | 98.7 |
| A235 | 97.8 | Shi.cat16-2 | 98.7 |
| ZJBH | 97.9 | A20 | 99.1 |
| BH1 | 98.0 | FR95-8958 | 99.1 |
| BH38 | 98.0 | Cat29-2022 | 99.2 |
| BH58 | 98.0 | Oit.cat70 | 99.3 |
| BH20 | 98.1 | Cat23-2023 | 99.4 |
| BH52 | 98.1 | HJ53 | 99.4 |
| BH54 | 98.1 | BH13 | 99.5 |
| BH56 | 98.1 | Tovarish1 | 99.5 |
| Zeus | 98.1 | Kyo.cat1 | 99.6 |
| BH27 | 98.2 | Kyo.cat31 | 99.6 |
| BH40 | 98.2 | PL18 | 99.6 |
| Cat1-2023 | 98.2 | 87-66 | 99.7 |
| Cat14-2023 | 98.2 | F1 | 99.7 |
| FR95-9015 | 98.2 | HJ54 | 99.7 |
| Osa.catM35-1 | 98.2 | PL257 | 99.7 |
| A76 | 98.3 | A71 | 99.8 |
| Cat15-2023 | 98.3 | H237-1 | 99.8 |
| Cat17-2023 | 98.3 | Oit.cat98 | 99.8 |
| Cat3-2022 | 98.3 | PL186 | 99.8 |
| Cat6-2023 | 98.3 | G5436 | 100.0 |

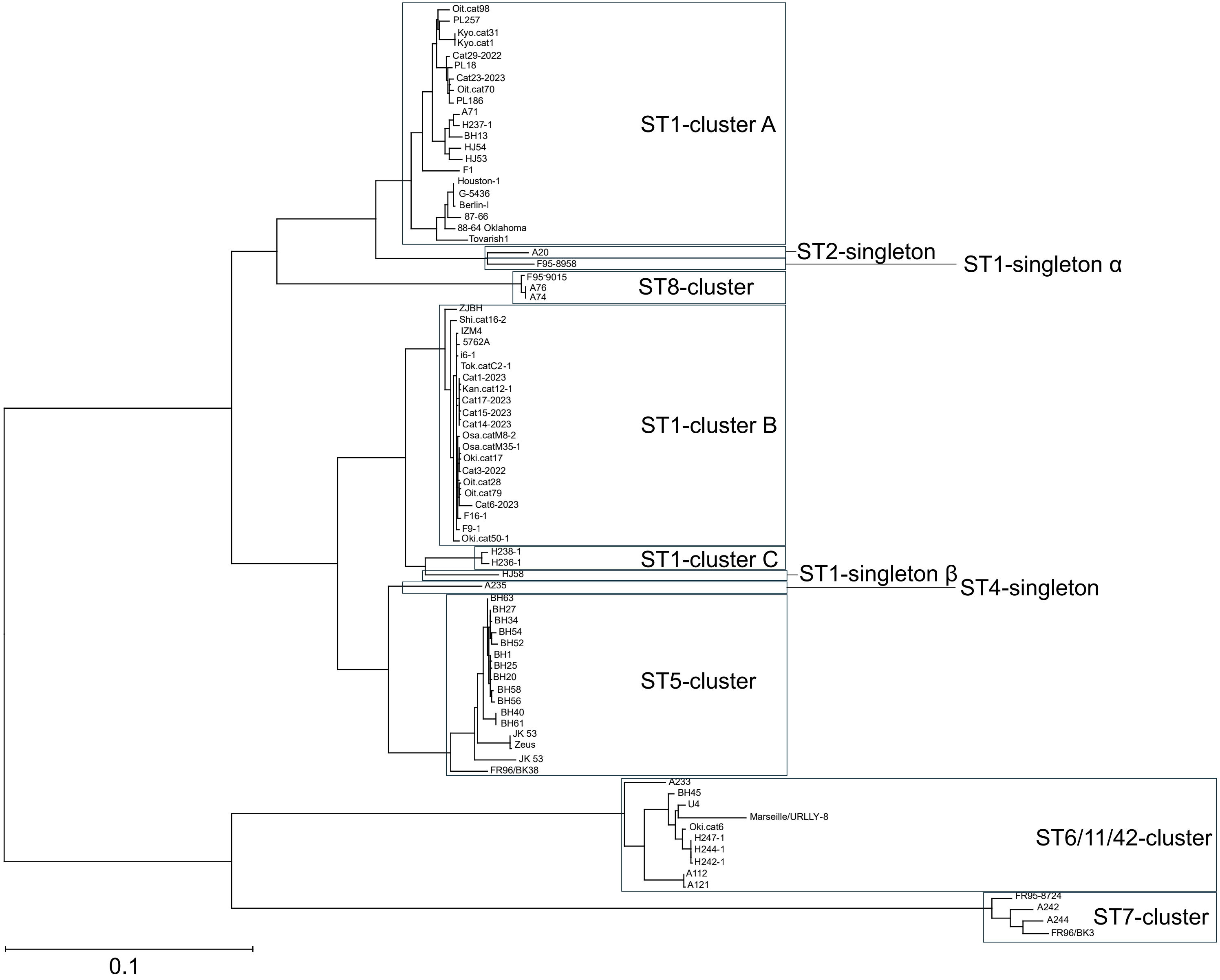
Oit.cat98
PL257
Kyo.cat31
Kyo.cat1
Cat29-2022
PL18
Cat23-2023
Oit.cat70
PL186
A71
H237-1
BH13
HJ54
HJ53
F1
Houston-1
G-5436
Berlin-I
87-66
88-64 Oklahoma
Tovarish1
ST1-cluster A
A20
F95-8958
ST2-singleton
ST1-singleton α
F95-9015
A76
A74
ST8-cluster
ZJBH
Shi.cat16-2
IZM4
5762A
i6-1
Tok.catC2-1
Cat1-2023
Kan.cat12-1
Cat17-2023
Cat15-2023
Cat14-2023
Osa.catM8-2
Osa.catM35-1
Oki.cat17
Cat3-2022
Oit.cat28
Oit.cat79
Cat6-2023
F16-1
F9-1
Oki.cat50-1
ST1-cluster B
H238-1
H236-1
ST1-cluster C
HJ58
ST1-singleton β
A235
ST4-singleton
BH63
BH27
BH34
BH54
BH52
BH1
BH25
BH20
BH58
BH56
BH40
BH61
JK 53
Zeus
JK 53
FR96/BK38
ST5-cluster
A233
BH45
U4
Marseille/URLLY-8
Oki.cat6
H247-1
H244-1
H242-1
A112
A121
ST6/11/42-cluster
FR95-8724
A242
A244
FR96/BK3
ST7-cluster
0.1

Figure S1
